# Transcription termination and antitermination are critical for the fitness and function of the integrative and conjugative element Tn*916*

**DOI:** 10.1101/2024.09.05.611371

**Authors:** Erika S. Wirachman, Alan D. Grossman

## Abstract

Premature expression of genes in mobile genetic elements can be detrimental to their bacterial hosts. Tn*916*, the founding member of a large family of integrative and conjugative elements (ICEs; aka conjugative transposons), confers tetracycline-resistance and is found in several Gram-positive bacterial species. We identified a transcription terminator near one end of Tn*916* that functions as an insulator that prevents expression of element genes when Tn*916* is integrated downstream from an active host promoter. The terminator blocked expression of Tn*916* genes needed for unwinding and rolling circle replication of the element DNA, and loss of the terminator caused a fitness defect for the host cells. Further, we identified an element-encoded antiterminator (named *canT* for conjugation-associated antitermination) that is essential for transcription of Tn*916* genes after excision of the element from the host chromosome. We found that the antiterminator is orientation-specific, functions with heterologous promoters and terminators, is processive and is most likely a *cis*-acting RNA. Insulating gene expression in conjugative elements that are integrated in the chromosome is likely a key feature of the interplay between mobile genetic elements and their hosts and appears to be critical for the function and evolution of the large family of Tn*916*-like elements.

**AUTHOR SUMMARY:** Horizontal gene transfer allows bacteria to rapidly acquire new traits that can enhance their adaptability to different conditions. Integrative and conjugative elements (ICEs) are mobile genetic elements that reside integrated in a bacterial chromosome and can transfer to another cell via cell-to-cell contact through the element-encoded secretion system. ICEs often confer beneficial traits to their hosts, including antibiotic resistances, symbiotic/pathogenic determinants, metabolic capabilities, and anti-phage defense systems. Tn*916*, the first-described ICE, was identified based on its ability to transfer tetracycline resistance in the pathogen *Enterococcus faecalis*, and is found in several Gram-positive species. Once transferred into a new cell, Tn*916* integrates into AT-rich sequences, sometimes downstream from a host promoter. We found that Tn*916* has a transcription terminator near one end of the element that blocks transcription from an upstream host promoter, thereby protecting cells from detrimental effects of premature expression of element genes. Further, we found that Tn*916* has a transcription antitermination system that is essential for expression of element genes after excision from the host chromosome. Our findings highlight the complex layers of transcriptional regulation that have evolved in ICEs, impacting host cell viability and the spread of the element.

## INTRODUCTION

Horizontal gene transfer helps drive microbial evolution, allowing bacteria to rapidly acquire new genes that can enhance their ability to thrive in different conditions. Integrative and conjugative elements (ICEs) are mobile genetic elements that normally reside integrated in a bacterial chromosome and can transfer to another cell via conjugation (Fig 1A). Once transferred, an ICE can integrate into the chromosome of the new host generating a transconjugant. ICEs often carry cargo genes that confer beneficial traits to their hosts, including antibiotic resistances, symbiotic/pathogenic determinants, metabolic capabilities, anti-phage defense systems, and many others [1–3]. While integrated in the host chromosome, cargo genes are often expressed, but most of the genes needed for the ICE lifecycle are not.

**Fig 1.**
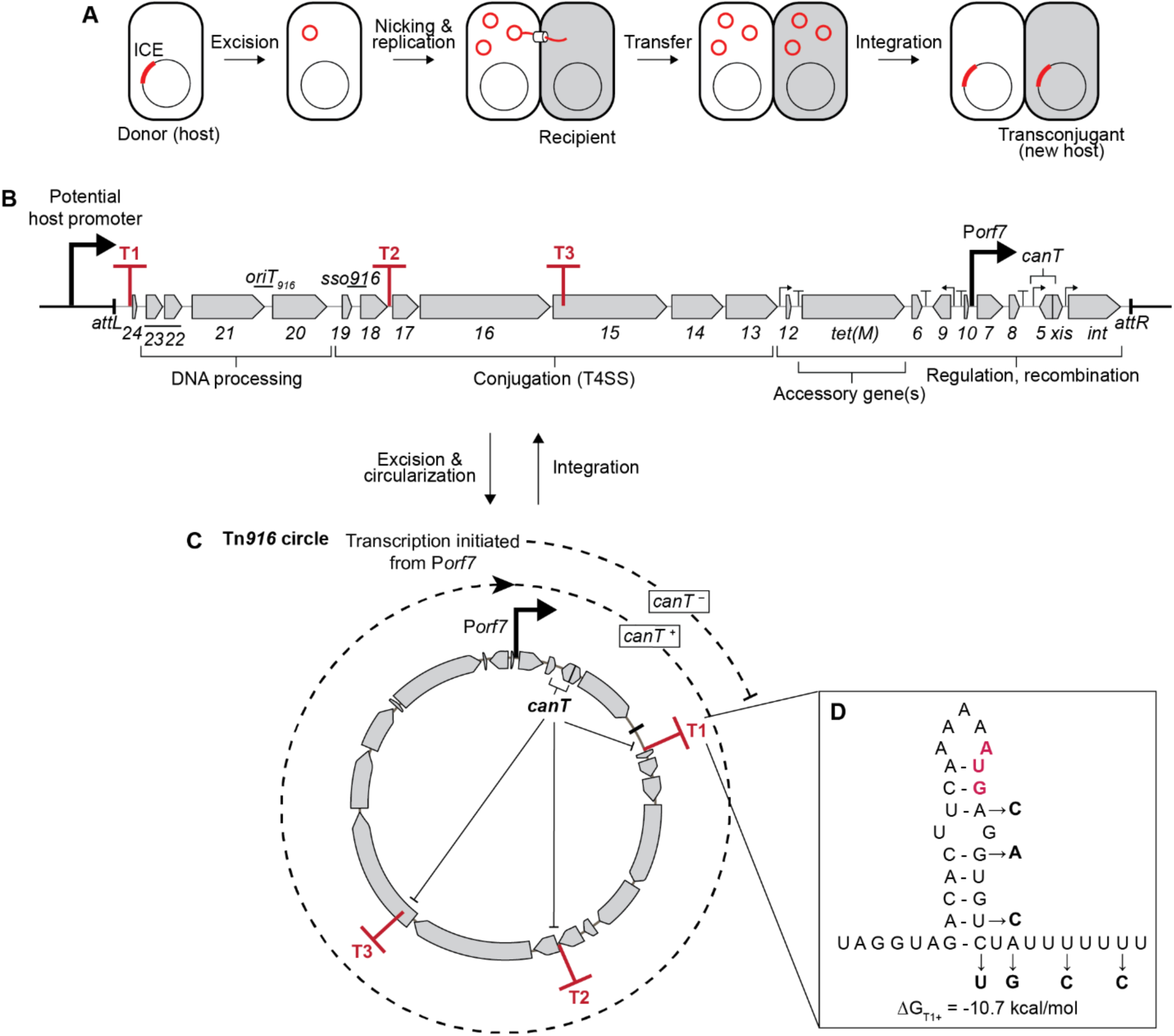
Genetic map of Tn*916* and its regulation. **A)** ICE life cycle. The chromosome is depicted as a circle with an ICE indicated in red. After excision, the circular double stranded ICE DNA (red) undergoes rolling circle replication. Transfer of linear ssDNA occurs to a recipient cell (shaded). Once transferred, the linear ssDNA is circularized and replicated to become dsDNA and then undergoes rolling circle replication and can be integrated to generate a stable transconjugant. **B)** Genetic map of Tn*916*. Gene names (numbers) are indicated below the corresponding genes. The ends of Tn*916* are indicated by black lines. The origin of transfer (*oriT_916_*) [66], single strand origin of replication (*sso916*) [65], and the antiterminator *canT* are indicated above the map. Three regions containing transcription terminators (T1, T2 consisting of T2a and T2b, T3) are indicated as red “T’s”. Functional modules of genes are indicated by brackets below the map. **C)** Tn*916* circle. Following excision, *canT* is now upstream from the element genes needed for DNA processing and conjugation and the *canT* RNA inhibits termination at T1, T2 and T3, thereby allowing transcription of the DNA processing and conjugation genes (*orf24*-*orf13*) from P*orf7*. **D)** RNA sequence of terminator T1. The nucleotide positions of the base of the terminator stem were determined by the ARNold web server [77,78]. The minimum free energy of folding ΔG (of the stem-loop) was calculated using the RNAfold web server [79]. Red, bolded AUG indicate the start codon of *orf24*. The changes in the T1 mutant are indicated and were made to preserve the potential ribosome binding site and amino acid sequence of *orf24*.

Tn*916* (∼18 kb) (Fig 1B), the first-described ICE, was identified based on its ability to transfer tetracycline resistance in the pathogen *Enterococcus faecalis* [4,5]. Tn*916* and its relatives are found in many Gram-positive species, including *Enterococcus*, *Streptococcus*, *Staphylococcus*, and *Clostridium* [4–12], and function quite well in *Bacillus subtilis* [13–18]. As with other ICEs, Tn*916* contains genes needed for its lifecycle: recombination (integration, excision); DNA processing (nicking, unwinding, and rolling circle replication); conjugation (a type IV secretion system); and regulation. When Tn*916* is integrated in the chromosome, its DNA processing and conjugation genes are not expressed, largely due to the absence of a promoter within the integrated element (Fig 1B). After excision (circularization) of the element, the DNA processing and conjugation genes are expressed from the promoter for *orf7* (P*orf7*) (Fig 1C) [19].

Tn*916* integrates into AT-rich genomic regions [20–23], which can be downstream from a host promoter. If Tn*916* genes are co-directional with a host promoter, then this promoter might drive expression of Tn*916* replication and conjugation genes (Fig 1B). Based on analogy to ICE*Bs1* from *B. subtilis* [24,25], we postulated that expression of these genes when the element is integrated would be detrimental to host cells, and that Tn*916* might have a mechanism to prevent this.

Here, we describe a transcription terminator (T1) near the left end of Tn*916* (Fig 1B) that is important for the fitness of host cells by functioning as an insulator to prevent transcription of element genes when Tn*916* is integrated downstream from a host promoter. We also discovered a site in Tn*916*, *canT* (conjugation-associated antiterminator) (Fig 1B and 1C), that prevents termination at element terminators and allows transcription of the DNA processing and conjugation genes, but only after excision of Tn*916* from the host chromosome. *canT* is essential for horizontal transfer of Tn*916*.

## RESULTS

### Identification of functional transcription terminators in Tn*916*

We identified four predicted intrinsic transcription terminators (T1, T2a, T2b, T3) between the left end through the conjugation genes of Tn*916* (Fig 1B). Terminator T1 is upstream of *orf24*, near the left end of Tn*916* (Fig 1B and 1D). T2a and T2b (indicated as T2), are adjacent to each other and downstream of *orf18* (Fig 1B and S1A Fig) and T3 is within *orf15* (Fig 1B and S1B Fig). Because of its location near the left end of Tn*916*, we focused on T1 as a potential insulator of element gene expression when Tn*916* is downstream from a host promoter.

To test its efficiency, we cloned T1 between the IPTG-inducible promoter Pspank and *lacZ* at an ectopic chromosomal locus, outside of Tn*916* (Fig 2A). In the presence of T1, expression of *lacZ* was reduced to <5% of that in the absence of T1 (Fig 2B). Mutations in T1 (T1^−^) that disrupted the stem and the U-tract of the terminator (Fig 1D) restored expression of *lacZ* (Fig 2B), indicating that T1 is a functional terminator with an efficiency of >95%.

**Fig 2.**
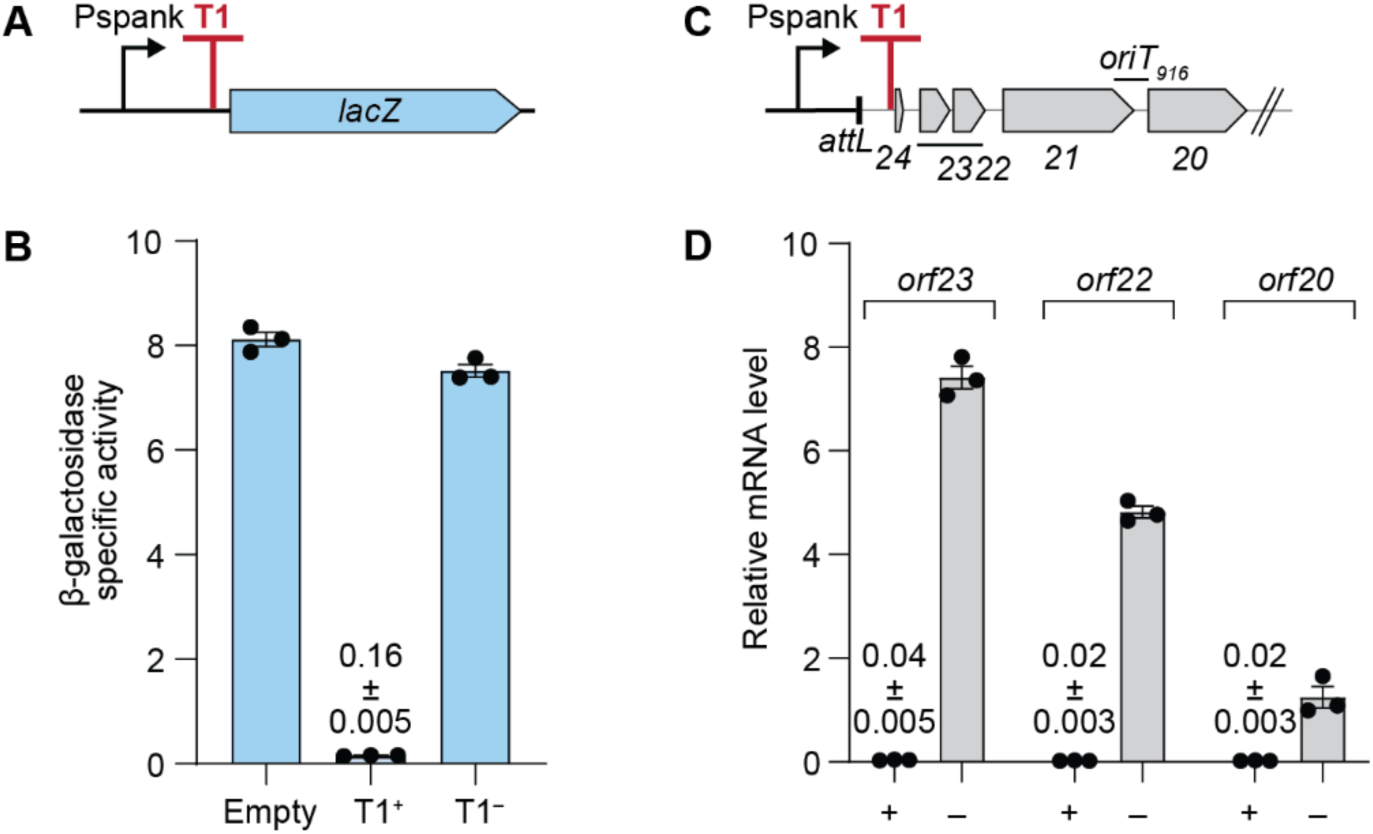
Terminator T1 of Tn*916* is functional within and outside of Tn*916*. **A)** Schematic of *lacZ* reporter construct with terminator T1 between Pspank and *lacZ*. **B)** β-galactosidase specific activities of the *lacZ* reporters with and without a functional terminator T1, either T1 was absent (Empty, ESW252), intact (T1^+^, ESW605), or mutated (T1^−^, ESW606), were measured two hours after induction of Pspank with IPTG. Data presented are averages from three independent experiments with error bars depicting standard error of the mean (mean ± SEM). **C)** Schematic of Pspank-Tn*916*. Full-length Tn*916* is present, but only *orf24* until *orf20* are indicated in the schematic. **D)** Relative mRNA levels of the indicated genes from Pspank-Tn*916* with an intact (T1+, ESW179) or mutant (T1–, ESW247) terminator T1 were measured one hour after induction of Pspank with IPTG. Data presented are from three independent experiments with error bars depicting standard error of the mean (mean ± SEM).

### T1 functions to insulate genes in Tn*916* from readthrough transcription from a promoter in the host chromosome

To test the ability of terminator T1 to insulate transcription of Tn*916* genes from an upstream promoter in the chromosome, we cloned Pspank upstream of Tn*916* with an intact or mutant T1 (Fig 2C) and measured the amount of mRNA of three Tn*916* genes (*orf23*, *orf22*, and *orf20*) located downstream of T1. The Pspank insertion prevented excision of the element (S2 Fig), thus, expression of Tn*916* genes was coming solely from the integrated element. After induction of Pspank (addition of IPTG), there were low levels of mRNA from the three genes in wild-type (T1^+^) Tn*916* (Fig 2D). In contrast, the T1 mutant (T1^−^) had elevated amounts of mRNA from each of the three genes (Fig 2D). Based on these results, we conclude that T1 is a functional terminator that greatly reduces transcription into Tn*916* from an upstream promoter in the chromosome.

### Predicting an antitermination mechanism in Tn*916*

Transcription of Tn*916* genes needed for conjugation is normally driven by P*orf7* after excision and circularization of the element from the chromosome [19] (Fig 1C). Because terminator T1 is between the promoter and conjugation genes, we postulated that there would be an antitermination mechanism to enable transcription of the Tn*916* genes essential for conjugative transfer. Previous work found that a transposon (Tn*5*) insertion in codon 41 (of 87 total codons) of the predicted gene *orf5* eliminated conjugation, leading to the inference that *orf5* might encode a protein essential for conjugation [26,27]. More recently, no transcripts from the putative *orf5* were detected [19,28]. Experiments described below demonstrate that the putative *orf5* protein product is not required for expression of Tn*916* genes or for conjugation. Rather, there is an overlapping site now called *canT* (conjugation-associated antitermination) that is essential for transcription antitermination and conjugation.

### The putative gene *orf5* does not encode a protein product necessary for conjugation

We made two different mutations that should prevent the production of an *orf5*-encoded protein. 1) We changed the predicted start codon from AUG to AUA [*orf5*(*M1I*)], which should prevent translation of *orf5* but preserve the amino acid sequence of the overlapping *xis*. 2) We changed codon 30 (of 83 total) to a stop codon [GCA to UAA, *orf5*(*A30\**)]. In both cases, the conjugation efficiency of Tn*916* was indistinguishable from that of an otherwise wild-type element (Fig 3A). These results indicate that the role of the putative *orf5* in conjugation is not as a protein-coding gene. More likely, the initial Tn*5* insertion in the putative *orf5* [26,27] disrupted a region that overlaps *orf5* and this region is required for conjugation.

**Fig 3.**
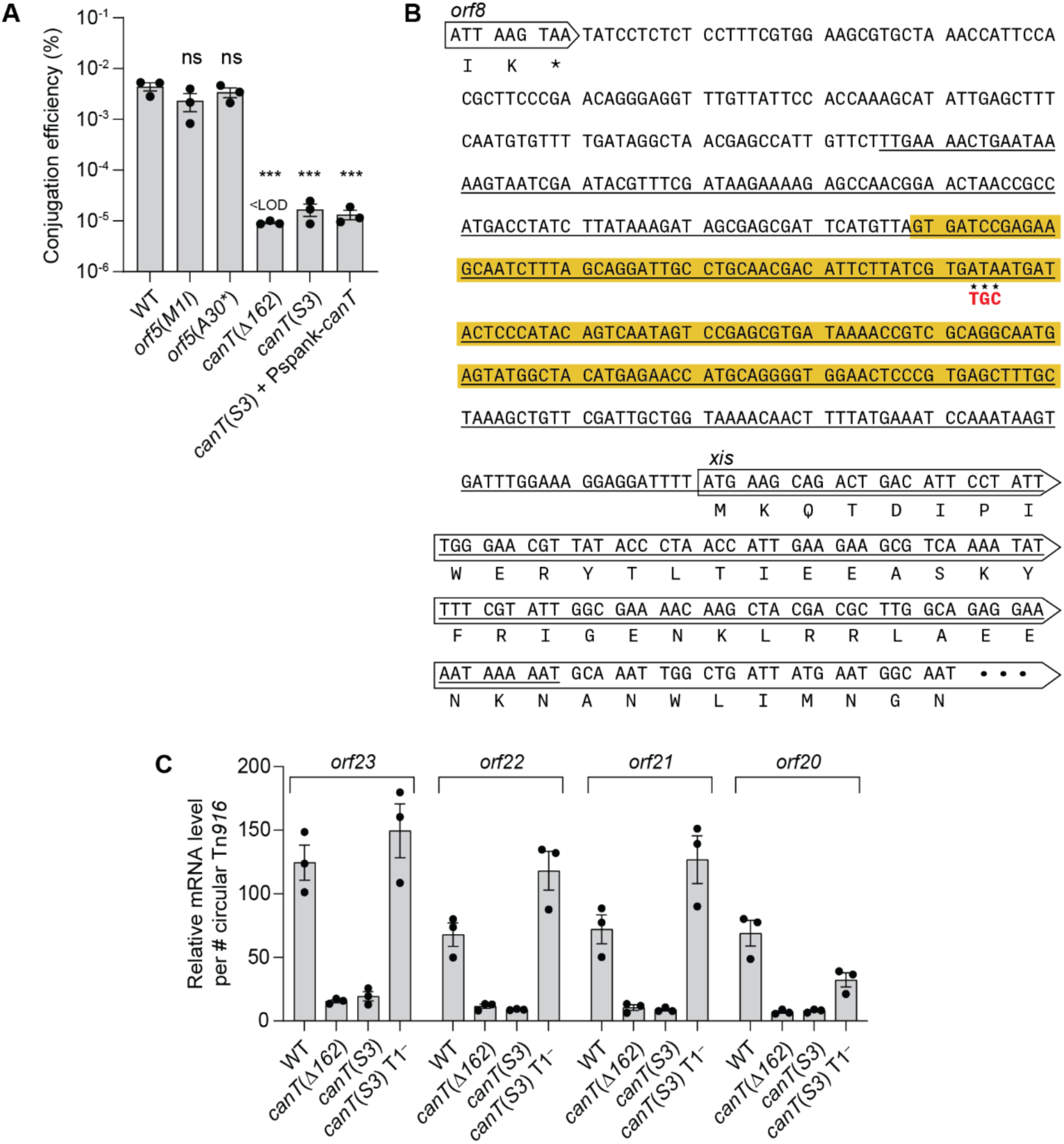
Phenotypes caused by mutations in *canT* and the putative *orf5*. **A)** Conjugation efficiencies. Donors contained wild-type Tn*916* (WT, CMJ253), Tn*916* with a mutation in the start codon of the putative *orf5* [*orf5*(*M1I*), ESW104], Tn*916* with a premature stop codon in the putative *orf5* [*orf5*(*A30\**), ESW420], Tn*916* with a deletion [*canT*(Δ*162*), ESW120] or 3 bp changes [*canT*(*S3*), ESW98] in *canT*, the Tn*916 canT*(*S3*) mutant with wild-type *canT* expressed in *trans* [*canT*(*S3*)+Pspank-*canT*, ESW771]. Conjugation efficiencies were calculated as the number of transconjugants per input donor. Data presented are from three independent experiments with error bars depicting standard error of the mean (mean ± SEM). All three independent mating assays of *canT*(Δ*162*) resulted in conjugation efficiencies that are below the limit of detection. P-values were calculated by an ordinary one-way ANOVA followed by Dunnett’s multiple comparisons test (*** *P*<0.001) using the GraphPad Prism version 10. Significance comparisons were made against the WT strain. **B)** Schematic of *canT* mutations. A region (452 bp) with *canT* is underlined. The region deleted (162 bp) in *canT*(Δ*162*) is highlighted in yellow. The changes in *canT*(*S3*) are indicated with stars and the sequence directly below. **C)** Relative mRNA levels normalized to the number of circular Tn*916* per cell. Tn*916* alleles include: wild-type (WT, CMJ253), *canT*(Δ*162*) (ESW120), *canT*(*S3*) (ESW98), *canT* and terminator T1 double mutant [*canT*(*S3*) T1^−^, ESW630]. Data presented are from three independent experiments with error bars depicting standard error of the mean (mean ± SEM).

### Identification of a region in Tn*916* that is required for transcription of conjugation genes

We found that the region between the 3’ end of *orf8* and *xis* was required for conjugation and expression of several Tn*916* genes. A deletion that removed 162 bp [*canT*(*Δ162*)] (Fig 3B, yellow-highlighted) and a 3 bp substitution (ATA to TGC) [*canT*(*3S*)] (Fig 3B) both caused an approximately 100-fold drop in conjugation (Fig 3A). In contrast to the effect on conjugation, the excision frequency of the mutants was not decreased (WT: 1.68 ± 0.06 %; *canT*(*Δ162*): 2.04 ± 0.05 %; *canT*(*3S*): 1.82 ± 0.06 %), indicating that the region altered in these mutants is normally required for conjugation but not excision, similar to the phenotypes described for the Tn*5* insertion in the putative *orf5* [26,27].

We also found that this region was required for expression of genes downstream from the terminator T1. We measured the amount of mRNA for *orf23, orf22, orf21,* and *orf20* per copy of circular Tn*916*, thereby normalizing for excision and copy number of the excised element. The amount of mRNA for these genes in the two different *canT* mutants was greatly reduced (Fig 3C), indicating that the normal function of *canT* is to enable expression of these genes. Together, these results show that *canT* affects expression of genes downstream from T1 in the Tn*916* circle, but is not required for excision.

If *canT* enabled expression of these genes by allowing transcription to read through T1 (as opposed, for example, to containing a strong promoter), then inactivation of T1 should restore gene expression in the absence of *canT*. Indeed, we found that loss of T1 restored expression of *orfs23, 22, 21,* and *20* in the *canT*(*3S*) mutant. Levels of mRNA from all four *orfs* were increased at least four-fold in the *canT*(*S3*) T1^−^ double mutant compared to the *canT*(*S3*) single mutant (Fig 3C). Based on these results, we conclude that *canT* functions as an antiterminator in Tn*916*, enabling transcription to read through terminator T1.

### *canT* is sufficient for antitermination and functions processively and in *cis*

#### *canT* functions as an antiterminator in the absence of other Tn*916* genes

We cloned a 452 bp fragment (Fig 3B, underlined) that contains wild-type or mutant *canT* between Pspank and T1 in the *lacZ* reporter described above (Fig 2A and 4A). Expression of *lacZ* with the *canT*(Δ*162*) and *canT*(*S3*) mutants was <5% of that with wild-type *canT* (Fig 4B). These results indicate that *canT* functions to allow transcription to read through T1, consistent with the results with an intact Tn*916*. Wild-type *canT* in the opposite (flipped) orientation did not allow expression of *lacZ* (Fig 4B), indicating that *canT* function is orientation-specific. We note that clones with wild-type *canT* in each orientation contain the putative *orf5*, which overlaps *canT*. In the construct with the opposite orientation, the putative *orf5* is oriented in the sense direction with Pspank and hence *orf5* should be transcribed. These findings reinforce the conclusions above that if *orf5* encodes a protein product, it is not involved in antitermination. Based on these results, we conclude that *canT* functions as an antiterminator, is orientation-specific, and no other Tn*916* genes are needed for *canT* function.

**Fig 4.**
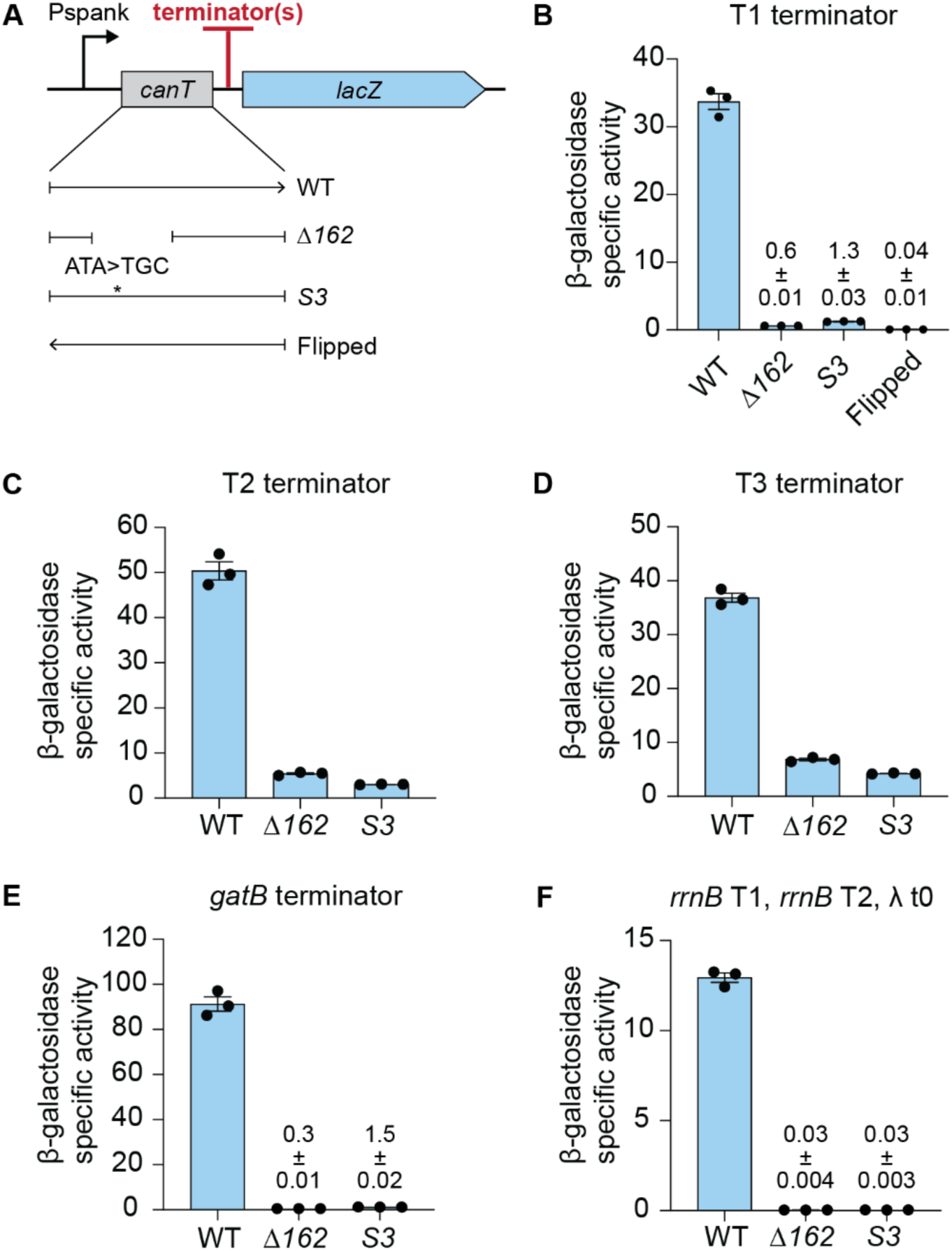
*canT* functions as an antiterminator without any other Tn*916* components. **A)** Schematic of *lacZ* reporter construct with *canT* alleles and terminator(s): T1 (**B**); T2 (**C**); T3 (**D**); *gatB* terminator (**E**); *rrnB* T1, *rrnB* T2, and λ t0 (**F**); between Pspank and *lacZ*. *canT* alleles included wild-type (WT), (Δ*162*), (*S3*), wild-type but in the opposite orientation (Flipped). **B-F)** β-galactosidase specific activities from the indicated strains following two hours induction of Pspank with IPTG. Data for each are from three independent experiments with error bars depicting standard error of the mean (mean ± SEM). **B)** T1 from Tn*916* with *canT* alleles: WT (ESW398), *Δ162* (ESW408)*, S3* (ESW407), Flipped (ESW440). **C)** T2 from Tn*916* with *canT* alleles: WT (ESW492), *Δ162* (ESW481)*, S3* (ESW623). **D)** T3 from Tn*916* with *canT* alleles: WT (ESW493), *Δ162* (ESW482)*, S3* (ESW626). **E)** Terminator from *gatB* with *canT* alleles WT (ESW421), *Δ162* (ESW473), *S3* (ESW449). **F)** Three terminators, T1 and T2 from *E. coli rrnB*, and t0 from phage lambda with *canT* alleles WT (ESW783), *Δ162* (ESW785), *S3* (ESW784).

#### *canT* functions as an antiterminator for T2 and T3 of Tn*916*

T2 and T3 were cloned individually into the *canT*-*lacZ* reporters in place of T1 (Fig 4A). Similar to results with T1, expression of *lacZ* was high in the presence of wild-type *canT* and much lower with the mutant *canT* (Fig 4C and 4D). The ability of *canT* to antiterminate T2 and T3 is consistent with the observation that genes located downstream of T2 and T3 are essential for conjugation and hence must be expressed in a wild-type Tn*916*.

#### *canT* functions as an antiterminator for heterologous terminators and is processive

We cloned the terminator located downstream of *B. subtilis gatB* into the *canT*-*lacZ* reporters (Fig 4A). Expression of *lacZ* was high with wild-type *canT* and quite low with the *canT* mutants (Fig 4E). We also cloned three terminators, the two from *E. coli rrnB* (*rrnB* T1, *rrnB* T2) and the phage lambda terminator t0, between *canT* and *lacZ* (Fig 4A). There was expression of *lacZ* with wild-type *canT*, but no detectable expression with the *canT* mutants (Fig 4F).

#### *canT* functions in *cis*

We wished to determine if there was a *trans*-acting factor that was disrupted in the *canT* mutant, or if *canT* functioned in *cis*. We found that a functional *canT* in the context of the reporter described above was unable to complement a Tn*916 canT* mutant. That is, there was no detectable conjugation of the Tn*916 canT* mutant in strains with a functional *canT* located elsewhere in the chromosome (Fig 3A). Based on the results above, we conclude that *canT* acts in *cis* to allow transcription to read through T1, T2, and T3 of Tn*916*, acts on heterologous terminators, and acts processively.

### Terminator T1 confers a fitness benefit to cells with Tn*916* integrated downstream from a strong promoter

We found that T1 of Tn*916* conferred a fitness benefit to the host when Tn*916* was downstream from and co-directional with a strong promoter in the host chromosome. Initially, we inserted the strong inducible promoter P*xis* from ICE*Bs1* upstream from Tn*916* (Fig 5A). P*xis* is repressed by ImmR and is derepressed following cleavage of ImmR by the protease ImmA which is activated by RapI [29–31]. We induced transcription from P*xis* by expressing *rapI* from a xylose-inducible promoter (Pxyl-*rapI*) (Methods). Transcription from P*xis* had little or no effect on the cell viability when Tn*916* contained a functional T1. That is, the number of viable cells four hours after derepression of P*xis* (the addition of xylose to induce Pxyl-*rapI*) was virtually the same as in cells grown similarly but without xylose (Fig 5B). In contrast, in cells with a mutant T1 (T1^-^), there was a two-fold reduction in CFUs after four hours of expression from P*xis* (+xylose) compared to no expression (no xylose) (Fig 5B). Based on these results, we conclude that terminator T1 is important for the fitness of host cells that contain Tn*916* downstream from a strong promoter.

**Fig 5.**
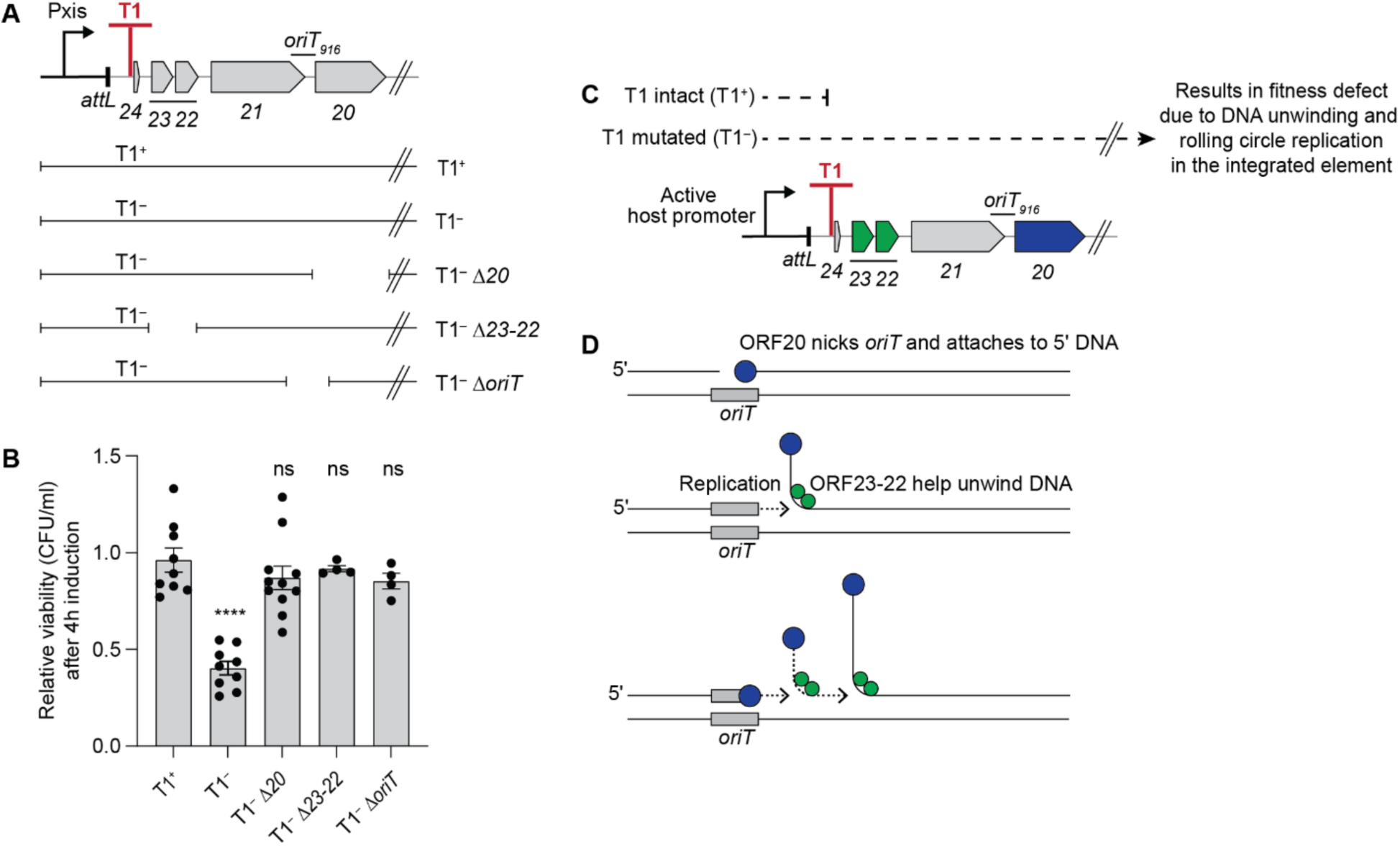
Rolling circle replication causes a fitness defect in Tn*916* insertions downstream of an active host promoter. **A)** Schematic of P*xis*-Tn*916* either with intact (T1^+^, ESW270) or mutant (T1^−^, ESW271) terminator T1, or with a mutant terminator combined with loss-of-function mutations in *orf20* (T1^−^ Δ*20*, ESW316), *orf23-22* (T1^−^ Δ*23-22*, ESW363), *oriT* (T1^−^ Δ*oriT*, ESW414). Full-length Tn*916* is present, but only *orf24* until *orf20* are indicated in the schematic. **B)** Relative viability of P*xis*-Tn*916* strains shown in (**A**) was measured four hours after induction of P*xis* and calculated as the number of CFUs following induction, divided by that in the uninduced culture grown in parallel (a value of “1” indicates there is no change in CFUs with induction). Data presented are from at least four independent experiments with error bars depicting standard error of the mean (mean ± SEM). P-values were calculated by an ordinary one-way ANOVA followed by Dunnett’s multiple comparisons test (**** *P*<0.0001) using the GraphPad Prism version 10. Significance comparisons were made against the T1*^+^* strain. **C)** Schematic of Tn*916* gene transcription when the element is inserted downstream of a host promoter. Genes necessary for DNA replication, including *orf23-22* (green) and *orf20* (blue), are not transcribed when terminator T1 is intact (T1^+^), but are transcribed when T1 is mutated (T1^−^). Expression of these genes from integrated Tn*916* leads to DNA unwinding and rolling circle replication occurring in the integrated element, which causes a fitness defect to the host cells. **D)** Cartoon of repeated rolling circle replication from the Tn*916 oriT* that is integrated in the chromosome. The relaxase ORF20 (blue circles) nicks the origin of transfer, *oriT* (gray bar), that also functions as an origin of replication [18] and is covalently attached to the 5’ end of the DNA. Replication extends (dotted line with arrow) from the free 3’-OH and regenerates a functional *oriT* that is a substrate for nicking by another molecule of the relaxase. ORF23-22 (green circles) act as helicase processivity factors, helping the host-encoded helicase, PcrA (not shown), unwind the DNA [18,32]. The rest of the replication machinery (not shown) is composed of host-encoded proteins. Cartoon is modified from (Menard and Grossman, 2013) [24].

By analogy to findings with ICE*Bs1* [24,25], we suspected that the decrease in CFUs of cells with the Tn*916* T1 mutant was due to autonomous rolling circle replication of Tn*916* that remained integrated in the chromosome. Tn*916* has three genes (*orf23, 22, 20*) and a site (*oriT*) that are required for DNA unwinding, autonomous rolling circle replication [18], and conjugation. *orf20* encodes the relaxase that nicks the element at the origin of transfer, *oriT* (that also functions as an origin of replication), to initiate DNA unwinding and rolling circle replication. *orf22* and *orf23* both encode helicase processivity factors that help the host-encoded helicase PcrA unwind the element DNA after nicking [18,32]. Deletions of *orf20*, *orf23-22*, or *oriT* all alleviated the fitness defect caused by expression from P*xis* into Tn*916* in the absence of a functional T1 (Fig 5A and 5B). Based on these results, we conclude that the terminator T1 in Tn*916* is important for preventing expression of the DNA unwinding and replication genes in Tn*916* when the element is downstream from a strong promoter and that loss of this terminator results in a fitness defect that is due to DNA unwinding and-or autonomous rolling circle replication of Tn*916* while it is in the host chromosome (Fig 5C and 5D).

### Isolating cells with Tn*916* integrated downstream of a host promoter

We found that T1 also contributed to host fitness when Tn*916* was integrated downstream from endogenous host promoters. To isolate Tn*916* insertions downstream from active host promoters, we cloned *lacZ* near the left end of Tn*916*, upstream of T1 (Tn*916-lacZ*) (Fig 6A) and used a strain with this element as a donor for conjugation. Transconjugants were selected on plates with tetracycline and X-gal and blue colonies were picked and verified for the presence of Tn*916* and expression of *lacZ*. Several strains were chosen for further analyses and the sites of integration were determined by arbitrary PCR and sequencing (Methods). Strains with an insertion in *sdpA*, *sunT*, *fadR*, and *adaB*-*ndhF* (Fig 6B) were chosen for further analyses. We removed *lacZ* from each insertion and then introduced mutations that inactivate terminator T1. As a control, we used an insertion in *yvgT-bdbC* in which the Tn*916* replication and conjugation genes were not co-directional with a host promoter (Fig 6B).

**Fig 6.**
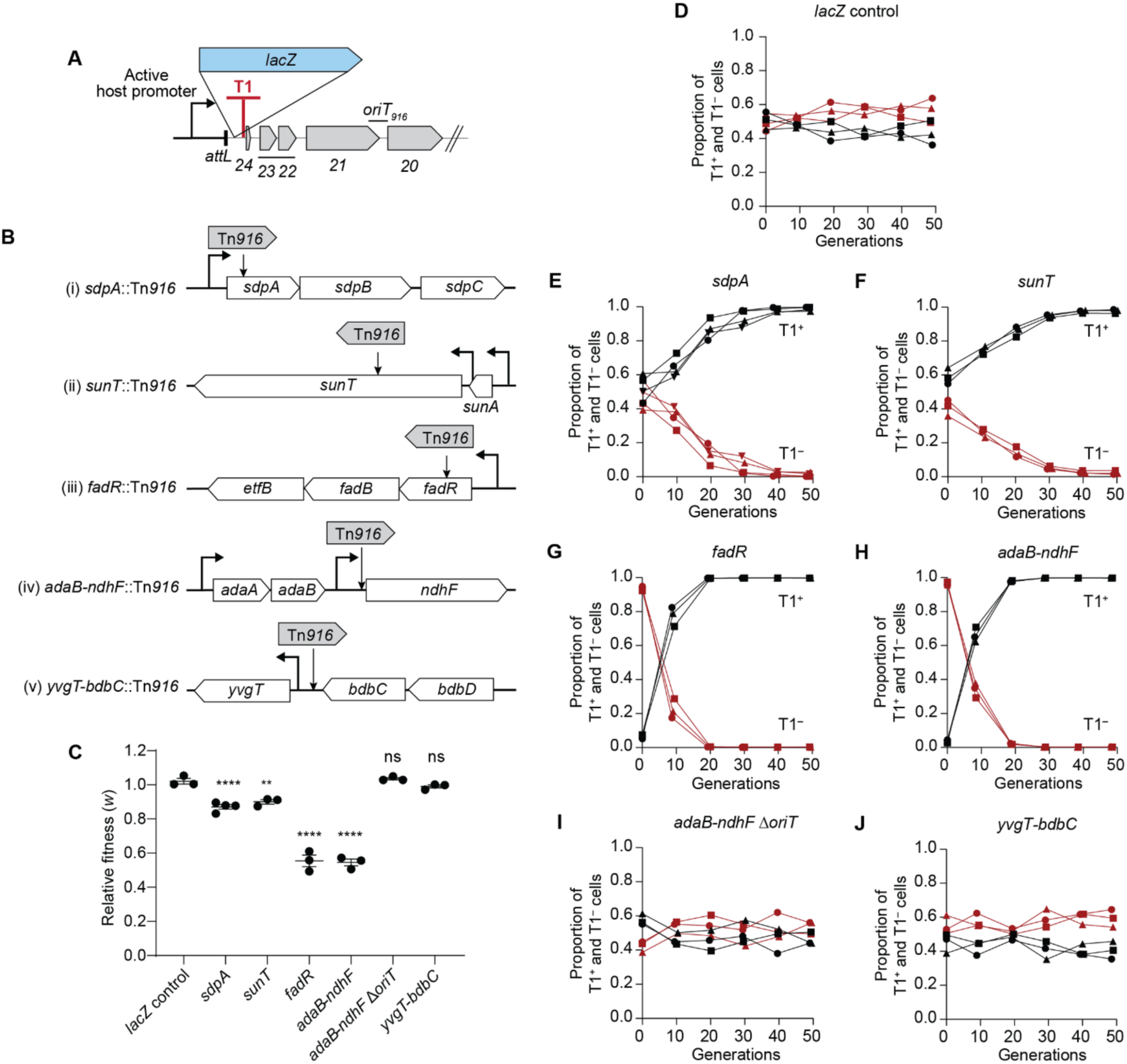
Terminator T1 confers a fitness benefit in Tn*916* insertions downstream of an active host promoter. **A)** Schematic of Tn*916*-*lacZ* strain (ESW261) used to isolate Tn*916* insertions that are downstream of host promoter. *lacZ* was cloned within Tn*916*, upstream of terminator T1. *lacZ* is expressed when Tn*916* integrates downstream of an active host promoter. **B)** Schematic and maps of the Tn*916* insertions used for pairwise competition assays. Tn*916* integrated at *sdpA* (i), *sunT* (ii), *fadR* (iii), and *adaB*-*ndhF* (iv) are located downstream of active promoters. Tn*916* at *yvgT-bdbC* (v) is oriented opposite the direction of transcription of the host genes. **C)** Fitness of T1^−^ cells relative to T1^+^ cells calculated as Malthusian parameters ratio of T1^−^ to T1^+^ (Methods). Data presented from at least three independent experiments with error bars depicting standard error of the mean (mean ± SEM). P-values were calculated by an ordinary one-way ANOVA followed by Dunnett’s multiple comparisons test (\*\**P*<0.01, \*\*\*\**P*<0.0001) using the GraphPad Prism version 10. Significance comparisons were made against the *lacZ* control. **D-J)** Population dynamics over the course of the pairwise competition assays. The initial ratio of *lacZ*^+^ and *lacZ*^−^ cells was ∼1:1 for (**D**). The initial ratio of T1^+^ to T1^−^ cells was ∼1:1 for (**E**,**F**,**I**,**J**) and ∼1:10 for (**G**,**H**). At least three independent experiments were done for each competition and individual data points are shown. The x-axis (generations) represents the number of population doublings. Replicate 1, circles; replicate 2, squares; replicate 3, triangles; replicate 4, inverted triangles. T1^+^, black; T1^−^, red. Competition assay results of *lacZ*^+^ and *lacZ*^−^ cells: (**D**) (ESW617 vs ESW616). Competition assay results of T1^+^ vs T1^−^ cells: (**E**) *sdpA* (ESW675 vs ESW693), (**F**) *sunT* (ESW677 vs ESW695), (**G**) *fadR* (ESW724 vs ESW725), (**H**) *adaB-ndhF* (ESW722 vs ESW723), and (**J**) *yvgT-bdbC* (ESW713 vs ESW714). (**I**) T1^+^ vs T1^−^ Δ*oriT* cells of Tn*916* integrated at *adaB-ndhF* (ESW722 vs ESW728).

### Fitness benefits of terminator T1 in Tn*916* insertions that are downstream from host promoters

To measure the fitness benefit conferred by T1, we did pairwise competition assays between strains with and without a functional T1 (T1^+^ versus T1^−^). The two strains were mixed in an appropriate ratio (∼1:1 or ∼1:10), spotted on LB agar for 24 hours, harvested, and then a dilution of the mix was spotted again on fresh LB agar and repeated for up to five days. The total cell number increased approximately 300- to 1,000-fold each 24-hour cycle, the equivalent of 8-10 cell doublings, and totaling approximately 50 cell doublings over five days. The proportion of each strain in the mixed population was determined every 24 hours, and the relative fitness (*w*) of T1^−^ to T1^+^ cells was calculated. Because *lacZ* had been removed from the Tn*916* insertions, we used *lacZ* located elsewhere in the chromosome as a fitness-neutral marker in T1^+^ cells (Fig 6C and 6D) to distinguish them from the T1^−^ cells.

We found that the loss of a functional T1 caused a fitness defect in the four different Tn*916* insertions that were downstream from a host promoter. The proportion of T1^−^ cells in the population dropped from ∼50% to ∼10% in ∼30 doublings for both *sdpA*::Tn*916* and *sunT*::Tn*916* (Fig 6E and 6F), with mean relative fitness of 0.87 and 0.90, respectively (Fig 6C). The proportion of T1^−^ cells in the population dropped from ∼90% to ∼1% in ∼20 doublings for both *fadR*::Tn*916* and *adaB-ndhF*::Tn*916* (Fig 6G and 6H), with mean relative fitness of 0.55 for both (Fig 6C). These results show that T1 can confer a fitness benefit to cells that contain Tn*916* downstream from and codirectional with a host promoter.

The fitness defect caused by the loss of T1 in the insertions was due to DNA unwinding and- or autonomous rolling circle replication. We deleted *oriT* from Tn*916* inserted in *adaB-ndhF*, thereby preventing both DNA unwinding and autonomous replication of Tn*916* and found that the proportion of T1^+^ and T1^−^ Δ*oriT* cells did not significantly change over the entire competition experiment (Fig 6I), indicating that the fitness defect caused by the loss of T1 was completely suppressed. The mean relative fitness was 1.04 (Fig 6C), in marked contrast to that of the element with a functional *oriT* (mean relative fitness of 0.55). These results are consistent with those described above for P*xis*-Tn*916* T1^−^ (Fig 5B).

The fitness effects caused by T1 were dependent on the presence of an upstream host promoter that was co-directional with Tn*916* genes. We used a Tn*916* insertion that is between *yvgT*-*bdbC* and oriented opposite the direction of transcription of the host genes (Fig 6B). There was no significant change in the proportion of cells with Tn*916* with T1^+^ versus T1^−^ during the entire competition (Fig 6J). The mean relative fitness was 0.99 (Fig 6C). Based on these results, we conclude that the terminator T1 of Tn*916* is important for the fitness of host cells when the element is downstream from and co-directional with an active host promoter, and that the primary function of T1 is to prevent autonomous rolling circle replication of Tn*916* that is in the host chromosome.

We anticipated that the fitness effects might be correlated with apparent promoter strengths. However, based on transcriptomics data [33], there was no obvious correlation between measured mRNA levels for each locus and the observed fitness. The transcriptomics data are from cells growing in liquid culture and not experiencing the complex growth patterns during colony formation on a solid surface, the conditions of our experiments. We believe that quantitatively different fitness effects are due to a combination of promoter strength, stability of the hybrid mRNAs (host gene and Tn*916* genes co-transcribed), and the ability of the hybrid mRNAs to be translated, all during complex growth conditions in the mixed communities on a solid surface.

## DISCUSSION

We found that Tn*916* has a transcription termination-antitermination system that is crucial for element function and host cell fitness. This system includes terminators T1, T2 (T2a + T2b), and T3 and the antiterminator *canT*. When Tn*916* integrates downstream from an active host promoter, T1 insulates expression of genes needed for unwinding and rolling circle replication of the element DNA and preserves host cell fitness. We suspect that the function of terminators T2 and T3 is to terminate spurious transcripts that might come from within the element, analogous to the proposed function of the terminators within the conjugation operon of the *B. subtilis* conjugative plasmid pLS20 [34].

After excision of Tn*916*, the promoter P*orf7* drives transcription of the DNA processing (nicking, unwinding, replication) and conjugation genes [19] and the terminators between P*orf7* and the end of the conjugation operon might be problematic for gene expression.. However, the *canT* antiterminator RNA allows transcription to read through element terminators processively, enabling expression of DNA processing and conjugation genes essential for conjugation once the element is in the circularized form (Fig 1C). This type of antitermination system appears to be widespread in the large family of Tn*916*-like elements.

### Factors involved in processive transcription antitermination

Processive transcription antitermination can involve protein and-or RNA factors, either acting independently or in combination. For example, the N and Q antitermination proteins of phage lambda bind to an RNA site (N) or a DNA site near a promoter (Q), and associate with RNA polymerase to cause antitermination [35–40]. The conjugative plasmid pLS20 encodes both the protein ConAn1 and the RNA *conAn2* that are used in antitermination of the long conjugation operon [34]. Antitermination systems in the lambdoid phage HK022 [41–44] and the *B. subtilis eps* operon [45] use RNA elements, *put* and EAR, respectively, for antitermination, without protein factors, although the possibility of host factor involvement has not been eliminated in the latter. Despite these variations, these factors are known or suspected to alter RNA polymerase such that it is no longer susceptible to most terminators [37,39,40,44,46–49]. We suspect that Tn*916 canT* RNA directly interacts with RNA polymerase to make it resistant to most terminators (Fig 7).

**Fig 7.**
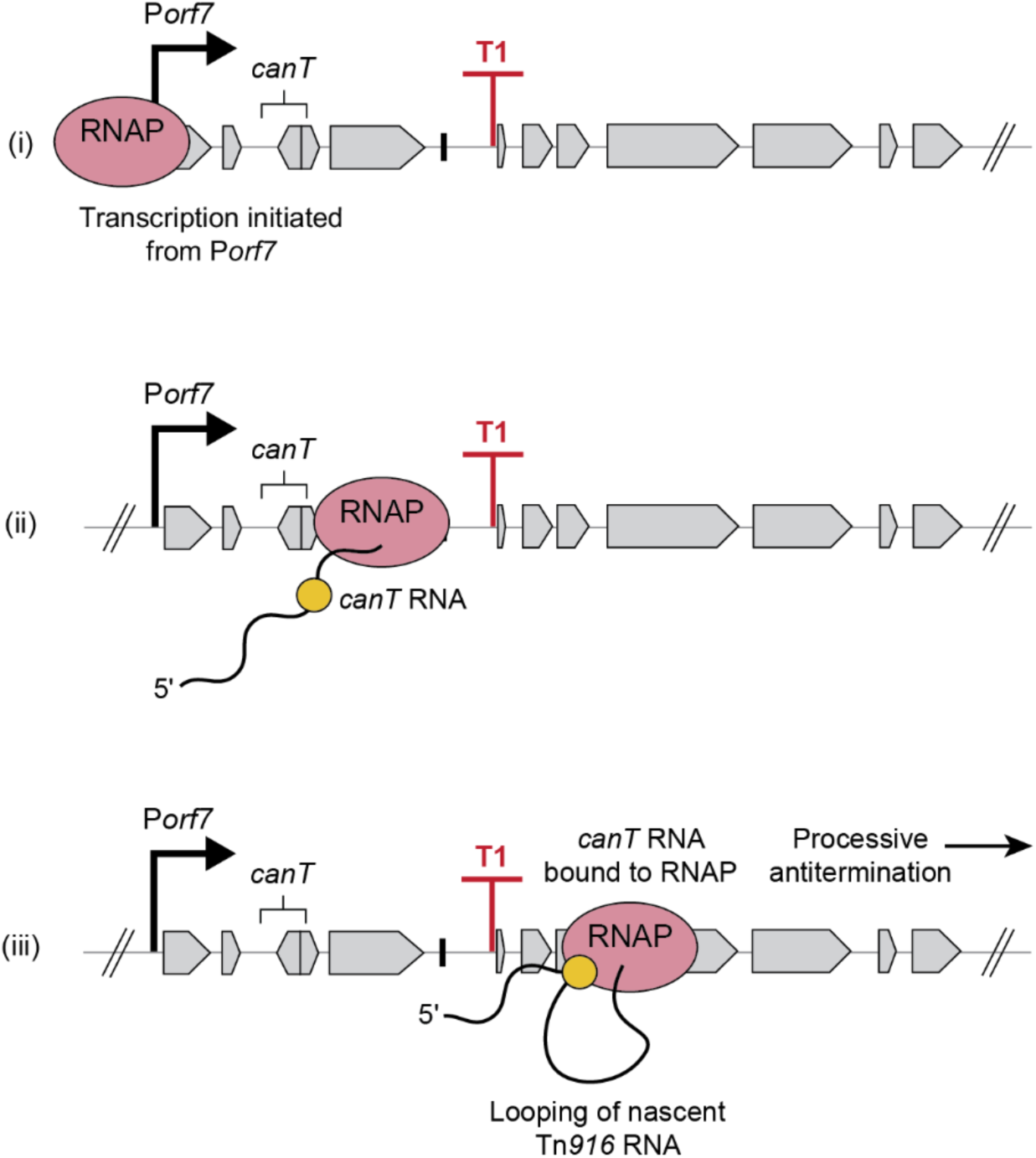
Cartoon for *canT*-mediated antitermination. (i) Post-excision, transcription by the RNA polymerase (pink) initiates at P*orf7* in the Tn*916* circle. (ii) The nascent RNA (wavy line) contains *canT* (yellow circle). (iii) *canT* RNA binds to the RNA polymerase, modifying it into a terminator-resistant form, allowing processive antitermination. *canT*-bound, modified RNA polymerase transcribes and bypasses terminator T1.

### Biological roles of processive transcription antitermination

Based on our findings and comparisons with other systems, Tn*916 canT* likely serves several roles in the biology of Tn*916*, including allowing complete transcription of the long DNA processing and conjugation operon despite the presence of internal terminators (discussed above), coupling element gene expression to excision, and enabling terminator T1 to function as an insulator to prevent element gene expression when Tn*916* is integrated downstream of a host promoter, thus, protecting host cells from detrimental effects of DNA unwinding and replication in the integrated element.

*canT* and P*orf7* couple element gene expression to excision. The DNA processing and conjugation operon is not expressed when Tn*916* is integrated due to the physical separation of the genes from the main promoter P*orf7* [19] (Fig 1B). Following excision, *canT* allows the transcript initiated from P*orf7* to bypass the internal terminators, leading to full expression of the operon in the Tn*916* extrachromosomal circle (Fig 1C). The use of antitermination to regulate timing of gene expression also occurs in the lambda phage, in which the N and Q proteins control the switch from immediate-early to delayed-early gene expression [35,37,39,40,50] and early to late gene expression [36–40,51], respectively.

### Mechanisms to prevent autonomous replication of integrated mobile genetic elements

ICEs have different mechanisms for regulating DNA nicking, unwinding and rolling circle replication. These mechanisms function to both couple autonomous replication with excision, and to stop new rounds of replication prior to or concomitant with integration. As described here, Tn*916* separates the main promoter and antiterminator from genes needed for DNA nicking, unwinding, and replication and uses a terminator upstream from those genes to prevent chromosomal promoters from reading into the element. This type of mechanism is enabled by the antitermination system in Tn*916*.

In contrast to Tn*916*, ICE*Bs1* uses a transcriptional repressor that controls expression of the excisionase (*xis*) and the DNA nicking, unwinding, and replication genes [29,52,53]. In this way, the DNA processing genes are expressed only when the excisionase is expressed. Further, repression and depletion of the excisionase is needed for integration, and this happens when *xis* is repressed, along with the DNA processing genes. This mechanism is similar to that used by some temperate phage, including lambda [54–57].

Our findings with Tn*916* highlight the intricate layers of transcriptional regulation in ICEs. We suspect that other ICEs may have similar termination-antitermination mechanisms and that this type of regulation is more widespread and may have a broader role in horizontal gene transfer than previously thought.

## METHODS

### Media and growth conditions

*B. subtilis* cells were grown shaking at 37°C in either LB medium or MOPS (morpholinepropanesulfonic acid)-buffered 1X S7_50_ defined minimal medium [58] containing 0.1% glutamate, required amino acids (40 μg/ml phenylalanine and 40 μg/ml tryptophan) and either glucose or arabinose (1% w/v) as a carbon source or on LB plates containing 1.5% agar. *Escherichia coli* cells were grown shaking at 37°C in LB medium for routine strain constructions. Where indicated, tetracycline (2.5 µg/ml) was added to Tn*916*-containing cells to increase gene expression and excision [19]. Antibiotics were otherwise used at the following concentrations: 5 μg/ml kanamycin, 10 μg/ml tetracycline, 100 μg/ml spectinomycin, 5 μg/ml chloramphenicol, and a combination of erythromycin at 0.5 µg/ml and lincomycin at 12.5 µg/ml to select for macrolide-lincosamide-streptogramin (*mls*) resistance. 5-bromo-4-chloro-3-indolyl β-D-galactopyranoside (X-gal) was used at a concentration of 120µg/ml.

Prior to sample collection for β-galactosidase assay, cells were grown as light lawns on 1.5% agar plates containing 1% w/v glucose, 0.1% w/v monopotassium glutamate, and 1X Spizizen’s salts [2 g/l (NH_4_)SO_4_, 14 g/l K_2_HPO_4_, 6 g/l KH_2_PO_4_, 1 g/l Na_3_-citrate·2H_2_O, and 0.2 g/l MgSO_4_·7H_2_O] [59]. Cells were resuspended from light lawns and grown at 37°C with shaking in 1X S7_50_ defined minimal glucose medium [58].

### Strains, alleles, and plasmids

*E. coli* strain AG1111 (MC1061 F’ *lacI*^q^ *lacZ*M15 Tn*10*) was used as a host for various plasmids. *B. subtilis* strains (Table 1), except BS49, were derived from JMA222, a derivative of JH642 (*trpC2 pheA1*) [60,61] that is cured of ICE*Bs1* [62]. *B. subtilis* strains were constructed by natural transformation [59] or conjugation as indicated. Key strains and newly reported alleles are summarized below.

**Table 1.**
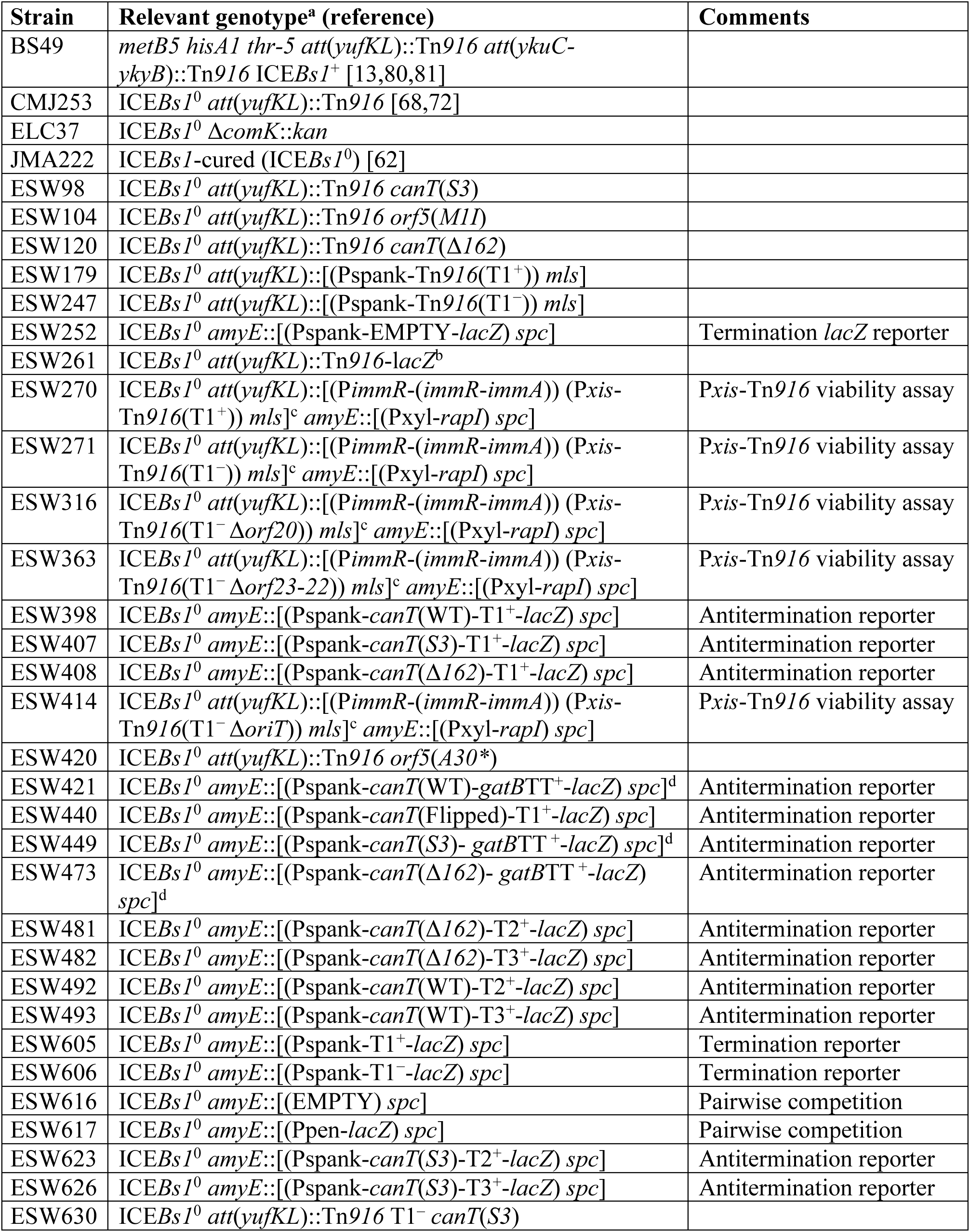
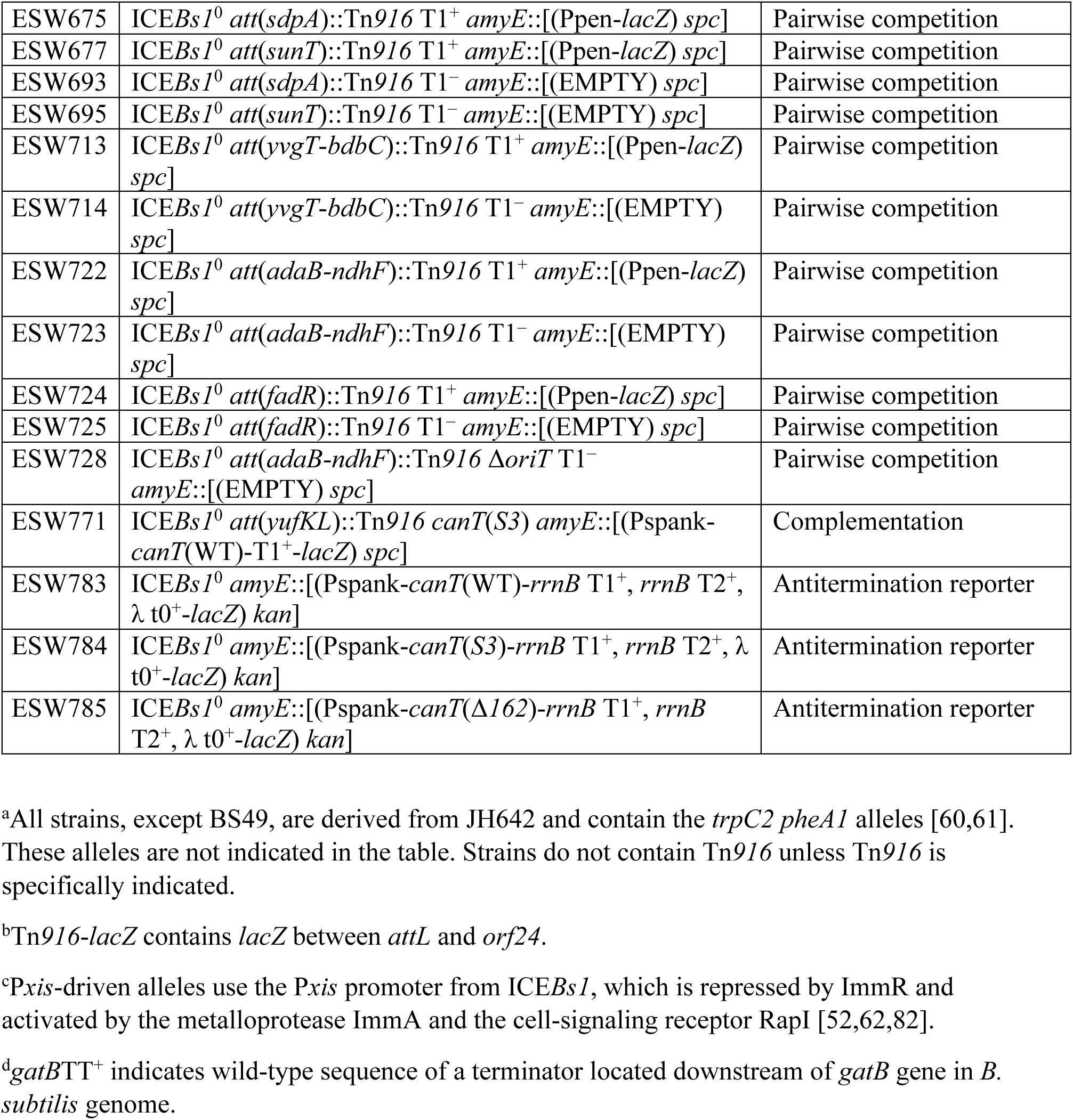
*B. subtilis* strains used.

*<u>ΔcomK</u>*<u>::*kan*</u> in ELC37 replaced most of the *comK* open reading frame from 47 bp upstream of *comK* to 19 bp upstream of its stop codon with the kanamycin resistance cassette (*kan*) from pGK67 [63]. The *kan* marker was fused with up- and downstream homology regions via isothermal assembly [64] and used for transformation.

#### Unmarked deletions and mutations in Tn*916*

Unmarked deletions and point mutations in Tn*916* were generated by a two-step allelic replacement approach. Briefly, DNA flanking the regions to be altered were amplified and inserted by isothermal assembly [64] into the EcoRI and BamHI sites of pCAL1422, a plasmid containing *E. coli lacZ* and *B. subtilis* chloramphenicol resistance cassette (*cat*), as previously described [32,65]. For deletion mutations, there was no DNA between the flanking regions. For point mutations, a fragment containing the desired mutations was assembled between the flanking regions. The resulting plasmids were used to transform the appropriate *B. subtilis* strains, selecting for integration of the plasmid into the chromosome (chloramphenicol-resistant) by single crossover recombination. Transformants were screened for loss of *lacZ* and checked by PCR and sequencing for the desired allele. Deletions and point mutations are described below.

*i) Genes and sites involved in replication.* Deletion of genes *orf23*-*22* (encoding helicase processivity factors) extends from immediately after the stop codon of *orf24* through the stop codon of *orf22*. Deletion of *orf20* (encoding the relaxase) fuses the first 90 codons of *orf20* with the *orf20* stop codon, deleting the intervening 306 codons and preserving *oriT* [66]. Deletion of *oriT* (Δ*oriT*) removes 5’-CCCCCCGTAT CTAACAGGGG GG-3’, starting from the nucleotide 41 downstream from the stop codon of *orf21*.

*ii) Mutations in the putative* orf5. *orf5*(*M1I*) (ESW104) changes the predicted start codon from AUG to AUA. This mutation preserves the amino acid sequence of the overlapping *xis*. *orf5*(*A30\**) (ESW420) changes codon 30 (of 83 total) from GCA (alanine) to UAA (stop).

*iii) Mutations in* canT. Deletion of *canT* [*canT*(*Δ162*), ESW120] removes 162 bp between *orf8* and *xis*, extending from 229 to 390 nucleotides downstream from the stop codon of *orf8*. *canT*(*S3*) (ESW98) contains a 3 bp substitution 5’-ATA-3’ to 5’-TGC-3’, starting from the nucleotide 283 downstream from the stop codon of *orf8*.

*iv) Mutations in the terminator T1.* Multiple base pair changes were made to inactivate terminator T1 (T1^−^), including changing three nucleotides in the predicted stem of the stem-loop, two nucleotides near the base of the stem, and two of the U’s that follow the stem in the predicted RNA secondary structure (Fig 1D). The mutations did not alter the ribosome binding site or amino acid sequence of *orf24*, which overlaps T1.

#### Pspank-Tn*916* and P*xis*-Tn*916*

Pspank-Tn*916* strains (T1^+^, ESW179; T1^−^, ESW247) and P*xis*-Tn*916* strains (T1^+^, ESW270; T1^−^, ESW271; T1^−^ Δ*orf20*, ESW316; T1^−^ Δ*orf23-22*, ESW363; T1^−^ Δ*oriT*, ESW414) were created by cloning, either Pspank or P*xis* promoter, and an MLS resistance cassette (*mls*) upstream of *att*(*yufKL*)::Tn*916* and its derivative mutants. DNA flanking the regions to be altered were generated by PCR and assembled flanking the promoter and *mls* via isothermal assembly [64]. DNA was transformed into *B. subtilis* selecting for resistance to MLS. Transcription from P*xis* was derepressed by overexpression of *rapI* under the control of a xylose-inducible promoter inserted at the non-essential locus *amyE* [*amyE*::(Pxyl-*rapI*) *spc*] [31]. RapI causes the protease ImmA to cleave ImmR, the repressor of P*xis*, thereby derepressing transcription from P*xis* [29,30].

#### Tn916-lacZ

Tn*916*-*lacZ* (ESW261) has *lacZ* at the left end of Tn*916*, upstream of T1 (Fig 6A) and was used to identify insertions that were downstream from an active promoter. Briefly, *lacZ* and a kanamycin resistance gene (*kan*) that was flanked by *lox* sites were inserted 29 bp upstream of *orf24* (upstream of T1) by isothermal assembly [64] and introduced into Tn*916*. The Cre recombinase, expressed from the temperature-sensitive plasmid, pDR244 [67], was then used to remove the *lox*-flanked *kan* marker by recombination. Strains were then cured of pDR244 by culturing them on LB agar at 42°C, as previously described [67,68]. *lacZ* is expressed when integrated Tn*916* is integrated downstream from an active promoter.

#### *lacZ* reporter for measuring transcription termination and antitermination

DNA fragments with possible terminators upstream from *lacZ* were cloned downstream of Pspank in the vector pDR110 (a gift from D. Rudner, integrates by double crossover at *amyE*; contains Pspank, *lacI*, *spc*). Briefly, *lacZ* was amplified by PCR from pCAL1422 [32] using primers that include either wild-type or mutant T1. The T1 sequence cloned included 6 bp and 15 bp directly upstream and downstream, respectively, of the base of the stem of the terminator hairpin. The PCR products were inserted by isothermal assembly [64] into pDR110, cut with NheI and HindIII and the resulting construct was integrated into *B. subtilis* at the *amyE* locus by recombination selecting for resistance to spectinomycin. Reporter strains contained: no terminator (ESW252); T1^+^ (ESW605); and the T1 mutant (T1^−^, ESW606).

Antitermination activity was measured using a reporter with a terminator between Pspank and *lacZ* and cloning additional DNA fragments containing the indicated *canT* alleles between Pspank and the indicated terminator. The 452 bp DNA fragment with wild-type *canT* extends from 126 to 577 bp downstream of the stop codon of *orf8*. The strategy for building each construct was similar to that described for the terminators above. Strains contained the *canT* alleles: WT, Δ*162, S3,* or flipped, followed by T1 (ESW398, ESW408, ESW407, ESW440, respectively); the WT, Δ*162, S3* alleles upstream of T2 (ESW492, ESW481, ESW623, respectively); upstream of T3, (ESW493, ESW482, ESW626, respectively); or upstream of the *gatB* terminator (ESW421, ESW473, ESW449).

Additionally, each of the three different *canT* alleles (WT, Δ*162, S3*) was cloned upstream from an array of three terminators, *E. coli rrnB* T1, *rrnB* T2, and the phage lambda λ T0, between Pspank and *lacZ* and integrated into the chromosome at *amyE*, essentially as described above, generating strains ESW783 (WT), ESW785 (Δ*162*), and ESW784 (*S3*).

### β-galactosidase assays

Cells were grown at 37°C in defined minimal medium with shaking. At OD_600_ ∼0.1, IPTG was added (1 mM final concentration) to induce transcription from Pspank and samples were taken two hours post-induction. Cells were permeabilized with 15 µl of toluene and β-galactosidase specific activity was determined [(ΔA420 per min per ml of culture per OD_600_ unit) × 1000] essentially as described [69] after pelleting cell debris.

### RT-qPCR to measure gene expression

For reverse transcription reactions, an aliquot of cells was harvested in ice-cold methanol (1:1 ratio) and pelleted. RNA was isolated using Qiagen RNeasy PLUS kit with 10 mg/ml lysozyme. iScript Supermix (Bio-Rad) was used for reverse transcriptase reactions to generate cDNA. Control reactions without reverse transcriptase were performed to assess the amount of DNA present in the RNA samples. RNA was degraded by adding 75% volume of 0.1 M NaOH, incubating at 70°C for 10 min, and neutralizing with an equal volume of 0.1 M HCl.

The relative amounts of cDNA were determined by qPCR using SSoAdvanced SYBR master mix and CFX96 Touch Real-Time PCR system (Bio-Rad). qPCR data were quantified using the standard curve method [70]. Standard curves for these reactions were generated using *B. subtilis* genomic DNA that contained wild-type Tn*916.* Primers used to quantify *xis* were oESW48 (5’-CTAACCATTG AAGAAGCGTC AAA-3’) and oESW49 (5’-ACGATTGCCA TTCATAATCA GC-3’). Primers used to quantify *orf23* were oELC436 (5’-GAAATGTTTT TCGCCAGCTT CAGC-3’) and oELC437 (5’-GCGAAGAATC AACGGACGGC-3’). Primers used to quantify *orf22* were oLW443 (5’-CTCTACGTCG TGAAGTGAGA ATCC-3’) and oLW444 (5’-TTGATAAGTT CCACCCGTGC G-3’). Primers used to quantify *orf21* were oELC438 (5’-CCTCACTACG TTTCATCATT TCTTCATAGA ATG-3’) and oELC439 (5’-GCTGACCTTG CGGACTTAGG-3’). Primers used to quantify *orf20* were oELC440 (5’-CTCTTTGCGT ACCAGTTCGC C-3’) and oELC441 (5’-GACCTTGCCA TTAACGATAA GACAGG-3’). Primers used to quantify the control locus *gyrA* were oMEA128 (5’-TGGAGCATTACCTTGACCATC-3’) and oMEA129 (5’-AGCTCTCGCTTCTGCTTTAC-3’). The relative mRNA levels of Tn*916* genes (as indicated by the Cq values measured by qPCR) were normalized to *gyrA* after subtracting the signal from control reaction without reverse transcriptase.

### qPCR to measure Tn*916* excision and circle copy number

qPCR was used to monitor excision (activation) and copy number of circular (excised) Tn*916*, essentially as described previously [18,71,72]. Briefly, cells containing Tn*916* were lysed (40 mg/ml lysozyme) and genomic DNA was prepared using the Qiagen DNeasy kit. qPCR was performed using SsoAdvanced SYBR master mix and the CFX96 Touch Real-Time PCR system (Bio-Rad). qPCR data were quantified using the Pfaffl method [73]. Standard curves for these qPCRs were generated using *B. subtilis* genomic DNA that contained an empty Tn*916* chromosomal attachment site (*att1*), an ectopic copy of the Tn*916* circle *att*Tn*916* junction inserted at *amyE*, and a copy of the nearby locus, *mrpG*.

Excision frequencies were calculated as the number of copies of the chromosomal site from which Tn*916* excised (*att1*) divided by the number of copies of *mrpG* (a nearby gene). The average number of copies of the circular Tn*916* per cell was calculated as the number of copies of *att*Tn*916* divided by the number of copies of *mrpG*. We used previously described primers [18]: oLW542 (5’-GCAATGCGAT TAATACAACG ATAC-3’) and oLW543 (5’-TCGAGCATTC CATCATACAT TC-3’) to amplify the empty chromosomal attachment site (*att1*), oLW526 (5’-AAACGTGAAG TATCTTCCTA CAG-3’) and oLW527 (5’-TCGTCGTATC AAAGCTCATT C -3’) were used to amplify the *att*Tn*916* junction in the circular Tn*916*, oLW544 (5’-CCTGCTTGGG ATTCTCTTTA TC-3’) and oLW545 (5’-GTCATCTTGC ACACTTCTCT C-3’) were used to amplify a region within the nearby gene *mrpG*.

### Growth and viability assays

P*xis*-Tn*916* strains were grown in defined minimal medium with 1% arabinose as a carbon source to early exponential phase. At an OD_600_ of 0.05, the cultures were split and xylose was added (1% final concentration) to one portion to induce transcription from Pxyl, thus expressing *rapI* (Pxyl-*rapI*), causing inactivation of ImmR, the repressor of P*xis* [29,30]. After four hours, the number of CFUs was determined in induced and non-induced cultures. “Relative viability” was calculated as the number of CFUs present in the induced culture divided by the number of CFUs present in the non-induced culture.

### Mating assay

Mating assays were performed essentially as described previously [62]. Briefly, donor strains containing Tn*916* (tetracycline-resistant) or derivatives were grown in LB medium to early exponential phase. At an OD_600_ ∼0.2, activation of Tn*916* was stimulated by addition of tetracycline (2.5 μg/ml final concentration). After one hour, donor strains were mixed in a 1:1 ratio with kanamycin-resistant recipient cells (ELC37) and 5 total OD units of cells were filtered. Mating filters were placed on a 1X Spizizen’s salts [59] 1.5% agar plate at 37 °C for one hour. Cells were then harvested off the filter and the number of CFUs of donors (tetracycline-resistant), recipients ELC37 (kanamycin-resistant), and transconjugants (tetracycline/kanamycin-resistant) were determined both pre- and post-mating. Conjugation efficiency is the percentage of transconjugants per donor (using the number of donors determined at the start of mating).

### Identification of Tn*916* insertions downstream from active host promoters

We used Tn*916-lacZ* to identify insertions downstream from host promoters. A donor containing Tn*916-lacZ* (ESW261) was crossed to a recipient without Tn*916* (ELC37) and cells were plated on LB agar containing tetracycline to select for transconjugants, kanamycin to kill donors (counterselection), and X-gal to screen for insertions downstream from an active host promoter, as indicated by blue colony color. Blue transconjugant colonies were re-streaked non-selectively on LB agar containing X-gal and subsequently confirmed to be resistant to tetracycline (due to Tn*916-lacZ*).

### Mapping Tn*916*-*lacZ* integration sites

Arbitrary PCR was used to map Tn*916*-*lacZ* integration sites, as previously described [74,75]. Briefly, blue, tetracycline-resistant colonies were used as a template in a PCR reaction containing arbitrary primers (oELC1003: 5’-GGCACGCGTC GACTAGTACN NNNNNNNNNT GATG-3’) paired with a primer to either the right (oELC1009: 5’-GACATGCTAA TATAGCCATG ACG-3’) or left (oELC1010: 5’-GAAGTATCTTTATATCTTCA CTTTTCAAGG-3’) end of Tn*916*. Purified PCR products were then amplified using oELC1004 (5’-GGCACGCGTC GACTAGTAC-3’) and oELC1011 (5’-GAACTATTAC

GCACATGCAA C-3’) for the right junction or oELC1012 (5’-CGTCGTATCA AAGCTCATTC ATAAG-3’) for the left junction. These PCR products were then sequenced with oELC1011 or oELC1012 and mapped to *B. subtilis* genome (Genbank accession number CP007800 [61]).

### Competition assays and fitness

#### Strains and growth

Tn*916* T1^+^ and T1^−^ strains were created by first replacing *lacZ* from Tn*916*-*lacZ* with *kan* flanked by *lox* sites and then removing *kan*. DNA fragments from upstream and downstream of *lacZ* (in Tn*916-lacZ*) were assembled by isothermal assembly [64] with either T1^+^ or T1^−^ and the *lox-*flanked *kan* and recombined into each Tn*916-lacZ* insertion by transformation and selection for resistance to kanamycin. The *lox*-flanked *kan* marker was then removed by Cre-mediated recombination (using pDR244 [67] as described above). Strains made include Tn*916* inserted in: *sdpA* (T1^+^, ESW675; T1^−^, ESW693), *sunT* (T1^+^, ESW677; T1^−^, ESW695), *fadR* (T1^+^, ESW724; T1^−^, ESW725), *adaB-ndhF* (T1^+^, ESW722; T1^−^, ESW723; T1^−^ Δ*oriT*, ESW728), *yvgT-bdbC* (T1^+^, ESW713; T1^−^, ESW714).

Because the insertions no longer contained *lacZ*, we were able to use a constitutively expressed *lacZ* (Ppen-*lacZ*) at *amyE* in the Tn*916* T1^+^ strains to distinguish them from the T1^−^ strains. *amyE*::[(Ppen-*lacZ*), *spc*] (ESW617) was made by PCR amplifying Ppen-*lacZ* from pCAL1422 [32] and inserting it into BamHI-BlpI-cut pAJW82 (a pDR110-derived vector that is lacking Pspank and *lacI*) by isothermal assembly [64], and then integrating it into *B. subtilis.* The cells with Tn*916* T1^−^ contained *amyE*::*spc* (ESW616) from pAJW82 with no insert. Competition experiments demonstrated that Ppen-*lacZ* did not affect fitness.

Strains with Tn*916* containing a wild-type or mutant T1 were grown in LB medium to early exponential phase. Strains were then mixed at the indicated ratio after adjusting their OD_600_ to ∼0.01 and 50 µl of each competition mixture was spotted on LB agar and grown for 24 hours (1 day) at 37°C. Every 24 hours, spots were resuspended in 1X Spizizen’s salts [59] and diluted to an OD_600_ of ∼0.01 and then 50 µl of this resuspension was spotted on fresh LB agar and grown for another 24 hours at 37°C. This was repeated for a total of 5 days. Under these conditions, cells progress through ∼10 doublings per growth cycle on LB agar. At generation (doubling) 0 (initial input) and every 24 hours, the ratio of the two strains in each mixture was determined by serially diluting and plating the appropriate dilution on LB agar containing X-gal and counting the number of *lacZ*^+^ (blue; Tn*916* T1^+^) and *lacZ*^−^ (white; Tn*916* T1^−^) colonies.

#### Fitness calculations

We calculated the fitness of the strain containing the mutant relative to that of the strain containing the wild-type terminator T1 using the equation derived from the Malthusian parameter estimate of fitness as described [76].

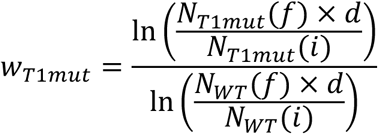

*w_T1mut_* is the fitness of the strain with Tn*916* T1^−^ relative to that with Tn*916* T1^+^. *N_T1mut_* and *N_WT_* are the numbers of CFU/ml of the strains with the mutant and wild-type T1, respectively. *i* indicates the initial number of CFU/ml of the indicated strain in the mixture and *f* indicates the CFU/ml after growth of the mixed strains. *d*, the dilution factor, is the fold-dilution of the cells from the start of the experiment (*i*) to the time (*f*) at which fitness was determined. Fitness was determined at a time when the population size was still changing (Fig 6D, 6E, 6F, 6G, 6H, 6I, and 6J). The dilution factor used to calculate fitness was ∼1,000 (two days of growth) for strains with insertions in *sdpA*, *sunT*, *adaB-ndhF* Δ*oriT*, and *yvgT-bdbC*, and the strains used for the *lacZ* control, and 1 (only one day of growth) for strains with insertions in *fadR* and *adaB-ndhF*.

Control competitions were performed to determine the fitness associated with the *amyE*::[(EMPTY) *spc*] marker used in Tn*916* T1^−^ cells (ESW616) relative to the *amyE*::[(Ppen-*lacZ*) *spc*] marker (ESW617) used in Tn*916* T1^+^ cells. The relative fitness was 1.02 ± 0.02 (mean ± SEM from three independent experiments), indicating that *lacZ* expression did not affect fitness of host cells (Fig 6C).

## ACKNOWLEDGEMENTS

We thank Mary E. Anderson for comments on the manuscript, Arielle J. Weinstein for plasmid pAJW82, Laurel D. Wright for previously constructing the Δ*orf23-22* and Δ*orf20* alleles used here, and Gabriel T. Vercelli for advice on fitness calculations.

---

**Fig S1.**
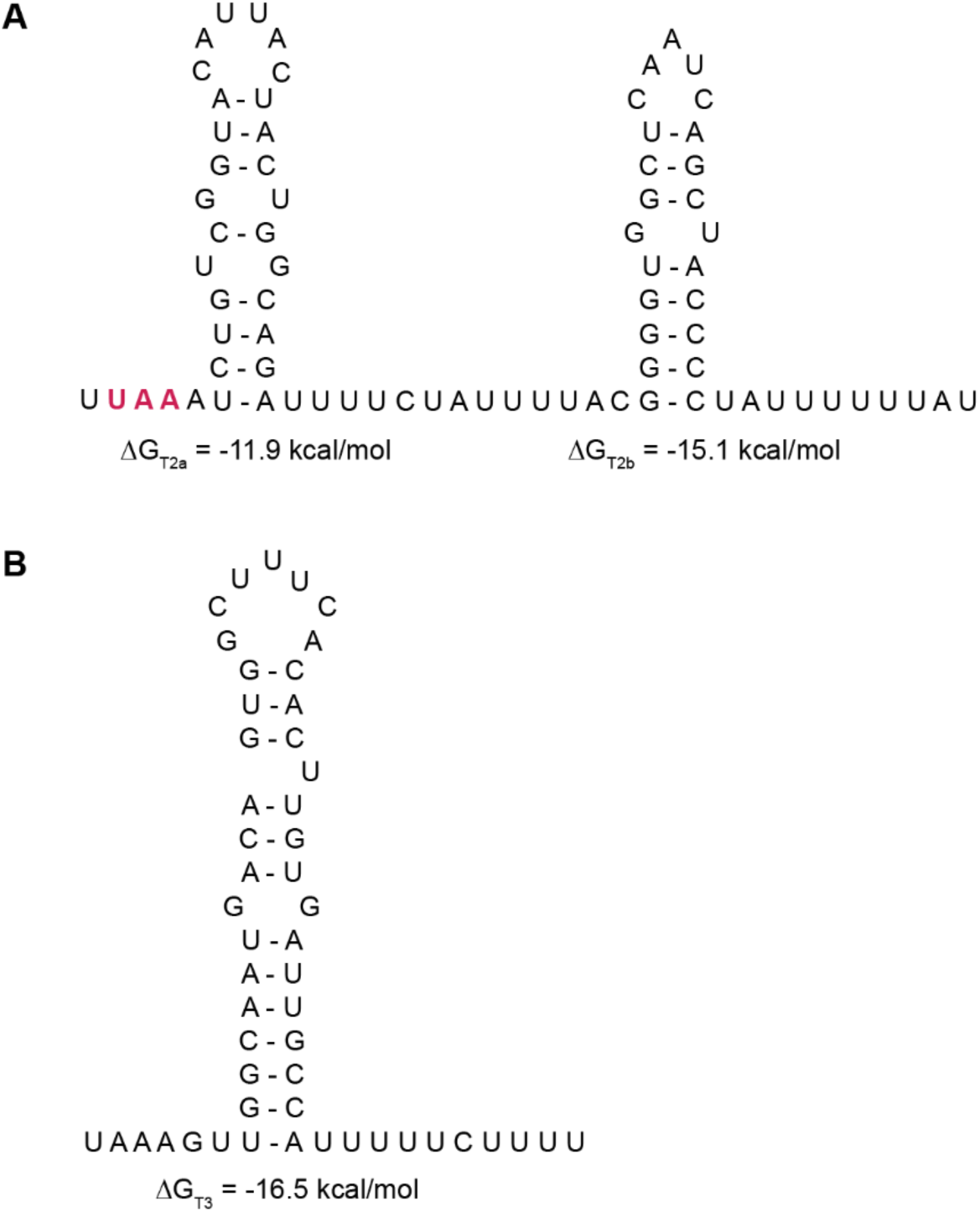
Terminator T2 and T3 of Tn*916*. The nucleotide positions of the base of the terminator stem were determined by the ARNold web server [77,78]. The minimum free energy of folding ΔG (of the stem-loop) was calculated using the RNAfold web server [79]. **A**) Terminator T2a and T2b. Red, bolded UAA indicate the stop codon of *orf18*. **B)** Terminator T3.

**Fig S2.**
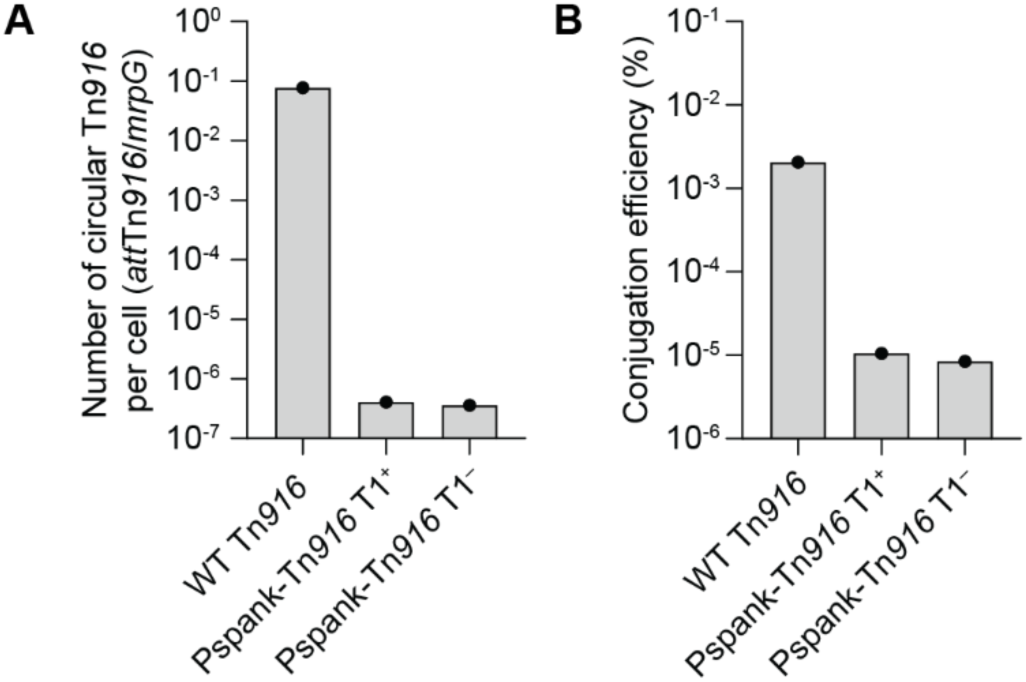
Pspank-Tn916 cannot excise and conjugate. **A)** Number of circular Tn*916* per cell (*att*Tn*916*/*mrpG*) and **B)** conjugation efficiencies of wild-type Tn*916* (CMJ253), Pspank-Tn*916* T1^+^ (ESW179), and Pspank-Tn*916* T1^−^ (ESW247). All strains were grown without tetracycline. Pspank-Tn*916* strains were grown continuously with 1 mM IPTG. Data presented are from one experiment. Mating assays of Pspank-Tn*916* T1^+^ and Pspank-Tn*916* T1^−^ resulted in conjugation efficiencies that are below the limit of detection.

